# Pupillometry as an objective measure of sustained attention in young and older listeners

**DOI:** 10.1101/579540

**Authors:** Sijia Zhao, Gabriela Bury, Alice Milne, Maria Chait

## Abstract

The ability to sustain attention on a task-relevant sound-source whilst avoiding distraction from other concurrent sounds is fundamental to listening in crowded environments. To isolate this aspect of hearing we designed a paradigm that continuously measured behavioural and pupillometry responses during 25-second-long trials in young (18-35 yo) and older (63-79 yo) participants. The auditory stimuli consisted of a number (1, 2 or 3) of concurrent, spectrally distinct tone streams. On each trial, participants detected brief silent gaps in one of the streams whilst resisting distraction from the others. Behavioural performance demonstrated increasing difficulty with time-on-task and with number/proximity of distractor streams. In young listeners (N=20), pupillometry revealed that pupil diameter (on the group and individual level) was dynamically modulated by instantaneous task difficulty such that periods where behavioural performance revealed a strain on sustained attention, were also accompanied by increased pupil diameter. Only trials on which participants performed successfully were included in the pupillometry analysis. Therefore, the observed effects reflect consequences of task demands as opposed to failure to attend.

In line with existing reports, we observed global changes to pupil dynamics in the older group, including decreased pupil diameter, a limited dilation range, and reduced temporal variability. However, despite these changes, the older group showed similar effects of attentive tracking to those observed in the younger listeners. Overall, our results demonstrate that pupillometry can be a reliable and time-sensitive measure of the effort associated with attentive tracking over long durations in both young and (with some caveats) older listeners.

## Introduction

The ability to sustain attention on a task-relevant stimulus whilst avoiding distraction from competing information is a fundamental perceptual challenge across sensory modalities. Arguably this is especially the case in hearing because of the dynamic nature of sound-objects. Listening in many natural environments (e.g. a busy train station, a loud restaurant, a noisy classroom) does not only depend on hearing acuity but also on the brain’s ability to focus and maintain attention on a specific sound (e.g. the announcement at the train station, a conversation in the restaurant, the teacher’s voice in the classroom) whilst resisting distraction from other concurrent sounds. Understanding ’attentive tracking’ is central to understanding the challenges faced by the brain during every-day listening and for addressing impairments in this ability. Indeed, diminished sustained attention capacity is hypothesized to underlie various disorders commonly associated with impaired listening, including Auditory Processing Disorder (APD; e.g. Moore et al., 2010, 2013), ADHD (Tucha et al., 2017), autism spectrum disorder (ASD; Corbett and Constantine, 2006) and dementia (Berardi et al., 2005; Calderon et al., 2001). Failure to maintain attention is also observed in hearing impaired individuals (Pichora-Fuller et al., 2016) and as a consequence of healthy ageing (Mishra et al., 2014; Petersen et al., 2017; Schoof and Rosen, 2014).

To successfully track a given source within a noisy scene, a listener must overcome challenges associated with *energetic masking* (i.e. extracting the information related to the target from the sound mixture) as well as challenges associated with selecting *and continuously following* the relevant source from within the background (Shinn-Cunningham and Best, 2008; Woods and McDermott, 2015). Most previous work has investigated listening in noisy environments using speech embedded in noise or in a mixture of other speakers, effectively confounding both aspects of tracking. However, it is likely that individual capacity to sustain attention is in itself a factor that will affect listening success. Here we sought to isolate and continuously monitor this aspect of auditory processing.

A large body of work demonstrates that sustained attention is not static but fluctuates over time and that these behavioural effects are associated with changes in connectivity along a distributed network of brain regions (Fortenbaugh et al., 2017; Langner and Eickhoff, 2013; Thomson et al., 2015). Emerging models postulate that lapses in sustained attention may arise from the weakening of executive processes over time, resulting in failure to effectively control resource allocation between the main task, distractor suppression, and mind wondering (Kurzban et al., 2013). To monitor sustained attention and determine how it is affected by increasing demands on distractor suppression, we designed a paradigm that isolates this facet of listening and measured behavioural and pupillometry responses during 25-second long trials.

Pupil dilation has long been used as a measure of effort (Beatty, 1982; Bradshaw, 1968; Cabestrero et al., 2009; Granholm and Steinhauer, 2004; Hjortkjær et al., 2018; Wel and Steenbergen, 2018) and is presently attracting considerable interest in the auditory modality because of evidence that pupil dilation can be used as an objective means with which to evaluate challenges to listening (McGarrigle et al., 2014; Peelle, 2018; Pichora-Fuller et al., 2016). The bulk of existing work has used pupillometry to evaluate listening effort associated with degraded speech (Koelewijn et al., 2012, 2014, 2015; Kuchinsky et al., 2014; Naylor et al., 2018; Ohlenforst et al., 2017; Wang et al., 2017; Wendt et al., 2016, 2017; Winn et al., 2015, 2018; Zekveld et al., 2010, 2011, 2018). As a result, these tasks inherently challenged both the ability to cope with energetic masking and the ability to sustain attention over time. Here we seek to specifically relate pupil dilation to the challenges of attentive tracking.

There is evidence to suggest that pupil dilation may be particularly correlated to the demands on sustained attention (Hopstaken et al, 2015; Sarter et al, 2001). Non-luminance mediated pupil dilation is at least partially driven by the release of NE (Norepinephrine, also Noradrenaline; Loewenfeld and Lowenstein, 1993) and ACh (Acetylcholine; see recent review Larsen and Waters, 2018). NE release has been consistently linked to arousal and sustained attention through its effects on modulating the response gain of cortical and thalamic neurons (Berridge and Waterhouse, 2003; Sara, 2009). ACh has been associated with activation in the anterior attention system and is hypothesized to play a role in controlling distraction (Berry et al., 2014; Demeter and Sarter, 2013; Kim et al., 2017; Sarter et al., 2006). We, therefore, expect that increased demands on sustained attention, including time-on-task and number of distractors. should be revealed in a time-specific manner in the pupil dilation pattern.

The auditory stimuli used in the present experiments are simple artificial ‘sound-scapes’ (Figure 1) that minimize the demands of segregation and isolate processes associated with object *selection*. ‘Scenes’ consist of a number (1, 2 or 3) of concurrent tone streams that model auditory sources. Each source is modulated at a unique rate; this contributes to the perceptual distinctiveness of each stream. The sources are widely set apart in frequency (always at least 6 ERB in Experiment 1; 2 ERB in Experiment 2) such that any effects are interpretable in the context of competition for processing resources rather than the increasing physical overlap between sources. On each trial, participants are instructed via a brief cue sound to attend to one of the streams. Attention is verified and quantified as performance on a gap detection task. Gaps occur in all streams but listeners are instructed to only respond to those in the target (‘Attended’) stream. The scenes are long (~25 seconds) and the task, therefore, requires listeners to maintain sustained attention over long durations and actively resist distraction from the other concurrent streams within the scene. As the scene size grows participants systematically struggle to resist the distractions, detecting fewer targets and making more false alarms (Figure 2). This demonstrates that this task models in a suitable way the competition for processing resources in crowded acoustic scenes.

**Figure 1.**
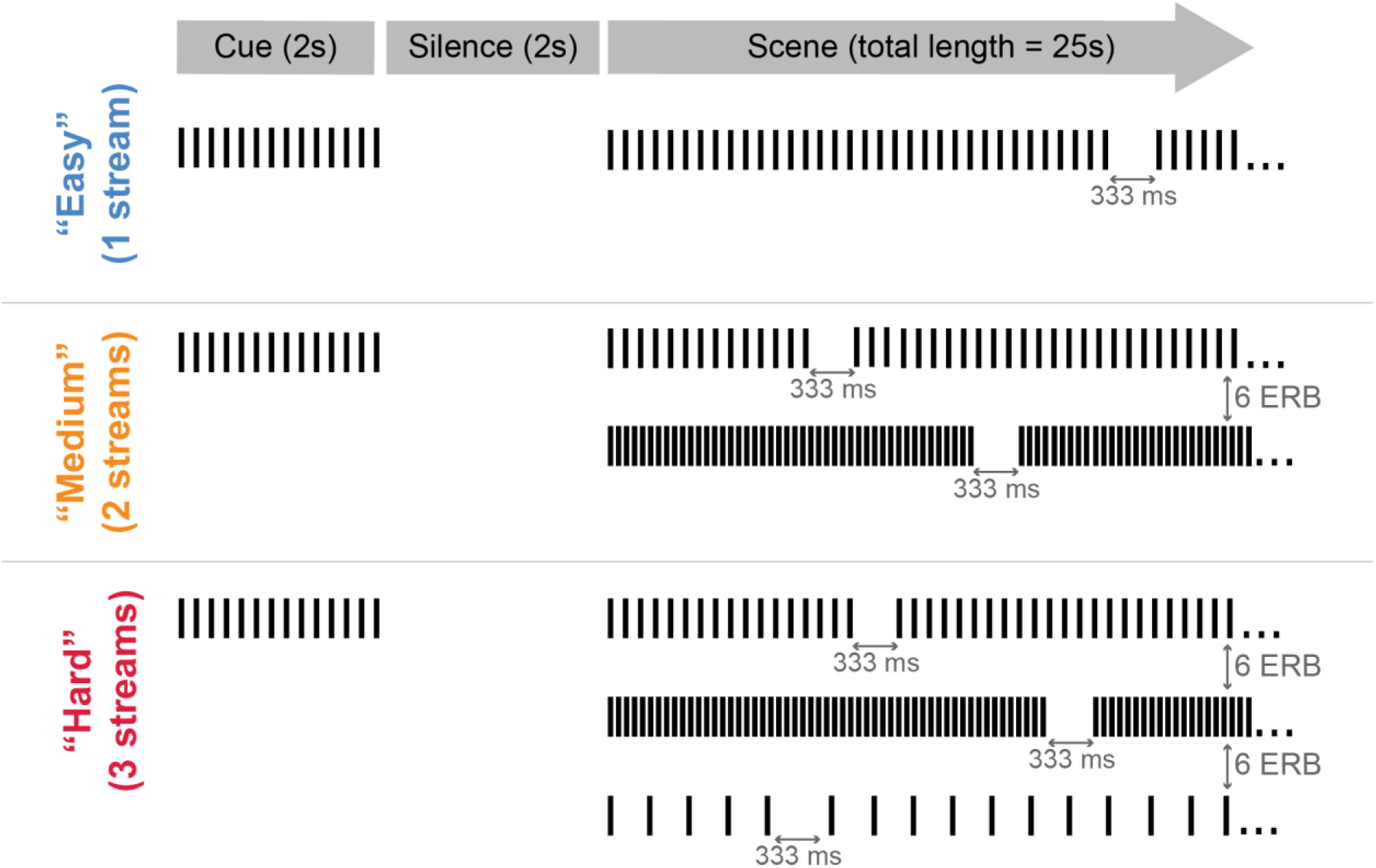
A schematic representation (not to scale) of the stimuli in Experiment 1. ‘Scenes’ consist of 1 (‘Easy’), 2 (‘Medium’) or 3 (‘Hard’) concurrent tone streams. Each source is amplitude modulated at a unique rate to increase distinctiveness. The sources are widely set apart in frequency (6 ERB). On each trial, participants are instructed (via a 2-second long cue sound) to attend to one of the streams (‘target’). Attention is verified and quantified as performance on a gap detection task. Gaps occur in all streams, but listeners are instructed to only respond to those in the target stream. The scenes are long (25 seconds) and as such the task requires listeners to maintain sustained attention over long durations and actively resist distraction from the other concurrent streams within the scene.

**Figure 2.**
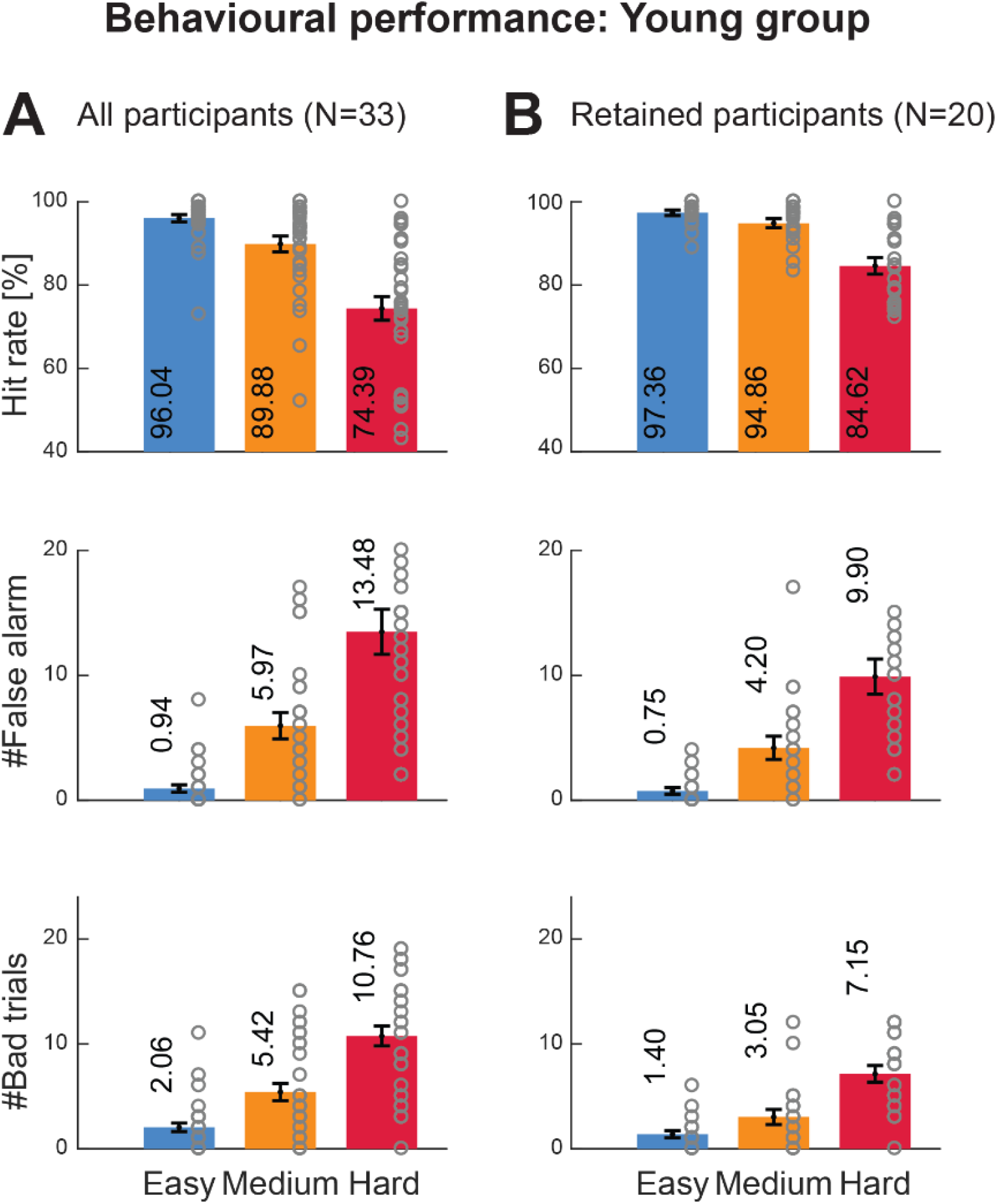
Behavioural performance of the Young group (Experiment 1). Performance measures were: hit rate, number of false alarms, number of bad trials. [A] Data from all participants (N=33). [B] Data from the participants retained for the pupillometry analysis (N=20). See ‘methods’ for retention criteria. Grey circles indicate individual data. The task conditions are labelled by difficulty: ‘Easy’ condition = 1 stream; ‘Medium’ condition = 2 streams; ‘Hard’ condition = 3 streams. All performance measures were significantly modulated by task difficulty. Error bar is ±1 SEM.

We address three questions: Firstly, we ask whether pupillometry can be a reliable and time-sensitive measure of the effort associated with attentive tracking over long durations similar to those over which listeners must maintain attention in ecologically relevant situations. Indeed, most previous work has used coarse pupil measures (peak dilation) and over relatively short intervals (most investigations have focused on the first 5 seconds; but see Hjortkjær et al., 2018). In contrast, we aim to measure instantaneous pupil diameter changes over a period of ~25 seconds.

A second challenge is related to isolating the effect of effort from other factors linked to task difficulty (McGarrigle et al., 2014). Manipulation of effort through varying task difficulty is intrinsically associated with reduced performance i.e. an increasing number of trials on which participants fail to accomplish the task. There is, therefore, a risk that pupil activity measured during those trials may reflect processes linked with the failure of attention and/or disengagement (for example, if the task is difficult participants might decide to ‘give up’ on a certain proportion of the trials). This is problematic because it is known that day dreaming is associated with pupil dilation (Franklin et al., 2013; Pelagatti et al., 2018). Indeed, Winn et al. (2015) compared pupil responses to correctly and incorrectly identified sentences and reported increased pupil dilation associated with failed trials, especially later in the trial (see their figure 1 and 5). This may be interpreted as indicating that the trials on which listeners failed were experienced as more difficult than the successful trials (see also Zekveld et al., 2010), but another interpretation may be that the increased dilation is related to disengagement rather than effort.

To address these concerns, we adopt a strict policy of analysing only successful trials. These are defined as trials on which all target gaps have been correctly identified and where the participant had at most one false positive. This allows us to focus on trials where resources were appropriately allocated, and distractors successfully ignored. Any differences observed between conditions will, therefore, reveal pure effects of task demands, not contaminated by failure to attend. Furthermore, we ensure that each condition and participant contribute an equal number of trials to the analysis (determined by the poorest performer on the hardest condition).

Finally, we ask whether pupillometry as a measure of effort to sustain attention is also applicable to older listeners. Attentive capacity is known to decline with age (e.g., Brosnan et al., 2018; Dørum et al., 2016; van der Leeuw et al., 2017; Lufi et al., 2015; Tu et al., 2018). An objective measure of sustained auditory attention would, therefore, be useful to quantify such difficulties and assess intervention outcomes. However, there are known changes to ocular physiology associated with healthy aging (Bitsios et al., 1996; Guillon et al., 2016; Tekin et al., 2018; Winn et al., 1994) that might limit the efficacy of pupillometry in this population (Piquado et al., 2010; Van Gerven et al., 2004).

Below, we report on two experiments run on young (18-31 year-old) and older (63-79 year-old) listeners. Overall, our results reveal that pupillometry can be a robust measure for attentive tracking in young listeners, supporting its potential usefulness as a screening measure and for evaluating various failures in sustained attention ability. There is also evidence that pupil measures may be used in older people but we identify a few cautionary issues.

## Experiment 1: Young listeners

### Methods

#### Participants

Thirty-three paid participants (21 females; mean age 22.9, range 18-31) took part in this study. All reported normal hearing and no history of neurological disorders. Experimental procedures were approved by the research ethics committee of University College London and written informed consent was obtained from each participant. Thirteen participants were excluded from the pupillometry analysis due to poor behavioural performance, leaving a subset of 20 participants (14 females, mean age 22.7, range 18-30).

#### Stimuli

Stimuli were 25-second-long artificial acoustics “scenes” that contained 1, 2 or 3 concurrent tone-pip sequences (“streams”). Each stream had a unique carrier frequency and modulation rate (AM). Carrier frequencies were selected from a pool of 18 ERB-spaced (Moore and Glasberg, 1983) values between 500 and 4000 Hz with the constraint that the separation between streams (in the 2- and 3-stream condition) was exactly 6 ERBs. AM rates were selected from a pool of 4 values 3, 7, 13 or 23 Hz. Tone pip duration was fixed at 30 ms (10 ms rise and fall). Together the unique combination of frequency and AM rate associated with each stream supported the perception of the scene as consisting of several concurrent, segregable “auditory objects”. In order to control for perceived loudness, the overall scene intensity was kept constant across scene-size conditions. As a consequence, individual stream intensity decreased with scene size.

Each stream contained between 2 and 3 silent gaps. These were created by removing the appropriate number of tones to generate a silent gap of around 333 ms (the minimum length of a gap in the 3Hz AM rate stream). Silent gaps could not occur within the first or last 2 seconds of a stream sequence or within 2 seconds of one another (including across streams). Participants were instructed to monitor one of the streams (“target”) for gaps whilst ignoring gaps in the distractor streams. The target stream was indicated by means of a 2000 ms cueing tone-pip sequences which preceded each trial. The scene was then presented following a 2000 ms silent gap (see Figure 1). In the 3-stream condition, the target stream was always the middle-frequency stream. To facilitate comparison across conditions, stimuli were created in triplets containing the same target stream across all 3 conditions. These were then presented in random order during the experimental session.

#### Procedure

Participants sat with their head fixed on a chinrest in front of a monitor (24-inch BENQ XL2420T with a resolution of 1920×1080 pixels and a refresh rate of 60 Hz) in a dimly lit and acoustically shielded room (IAC triple walled sound-attenuating booth). They were instructed to continuously fixate on a black cross presented at the centre of the screen against a grey background whilst monitoring the cued target stream for gaps. They were to respond (button press) as quickly as possible when a gap was detected whilst ignoring gaps in the distractor streams. Visual feedback (number of misses and false alarms) was presented for 1500 ms at the end of each trial.

Stimuli were presented in random order, such that on each trial the specific condition was unpredictable until scene onset. Sounds were delivered diotically to the participants’ ears with Sennheiser HD558 headphones (Sennheiser, Germany) via a Creative Sound Blaster X-Fi sound card (Creative Technology, Ltd.) at a comfortable listening level self-adjusted by each participant. Stimulus presentation and response recording were controlled with the Psychtoolbox package (Psychophysics Toolbox Version 3; Brainard, 1997) on MATLAB (The MathWorks, Inc.).

The entire experimental session lasted approximately 2 hours. Participants first completed a short practice block followed by 6 experimental blocks comprised of 12 trials each (~6.5min, 4 trials per condition). In total, 72 trials (24 trials per condition) were presented in a random order for each participant.

#### Analysis of behavioural data

Key presses occurring within 0.3 s of a previous keypress were considered to be accidental and removed from the analysis. A keypress was classified as a **hit** if it occurred 0.3 to 1.5 seconds following a target gap. **Hit rate (HR)** was computed for each subject, in each condition, as the ratio between detected vs. presented gaps in the target stream. All key presses that were not classified as a hit were classified as **false alarms (FA)**. These were summed and averaged across trials as a measure of distractibility. As mentioned above, only trials on which participants performed well were included in the pupillometry analysis. “Successful trials” were those where all the target gaps were correctly detected (100% hits) and which included at most one FA. All other trials were classified as “**bad trials**” and removed from the analysis. These 3 measures: HR, #FA and #bad-trials are plotted as measures of performance in Figures 2, 3, 4 and 5. Note that FA is quantified as a count (and not as a rate). This is because false responses can happen at any time during the trial.

**Figure 3.**
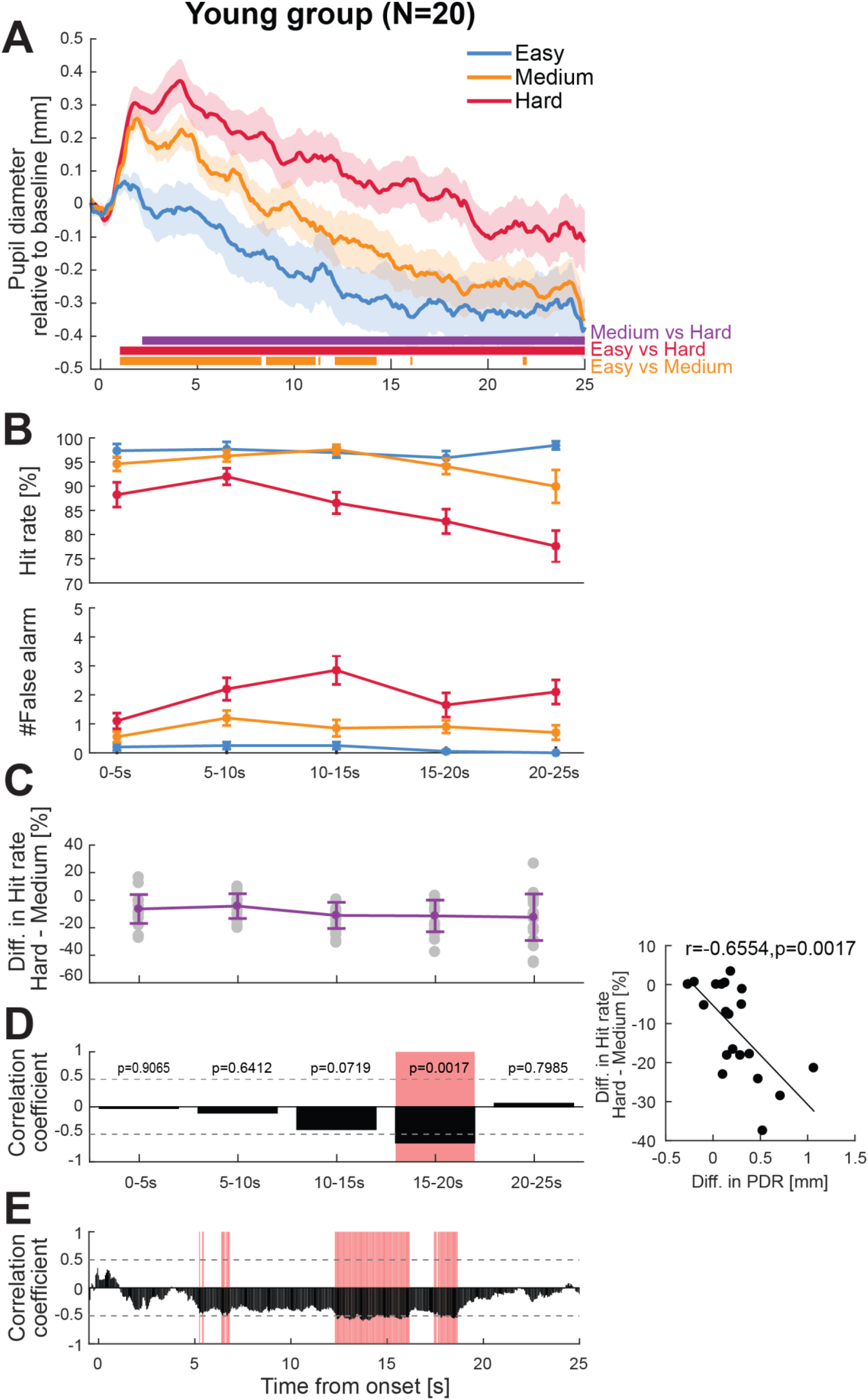
The pupil dilation response reflects effort to sustain attention. [A] Pupil dilation results from the young group (N=20). The solid lines represent the average pupil diameter as a function of time relative to the baseline (500 ms pre-onset). The shaded area shows ±1 SEM. Colour-coded horizontal lines at graph bottom indicate time intervals where bootstrap statistics confirmed significant differences between each pair of conditions. [B] Time-binned behavioural performance. Error bars are ±1 SEM. [C] Time-binned Hit Rate difference between the ‘Hard’ and ‘Medium’ conditions. Error bars are ±1 standard deviation. Grey dots represent individual data. [D] Correlation between PDR and HR for each time bin. Within each time-bin average PDR difference between the ‘Hard’ and ‘Medium’ conditions is correlated with the corresponding HR difference (as in [C]). Black bars indicate Spearman correlation coefficients at each time bin. Red shaded areas indicate time interval where a significant correlation (Bonferroni corrected) was observed. Plotted on the right-hand side is the correlation in the 15-20 sec time-bin. Each dot represents data from a single subject. [E] Correlation between PDR (‘Hard’ – ‘Medium’ condition) and behavioural performance (hit rate difference between the ‘Hard’ and ‘Medium’ conditions) on an individual subject level. Black bars indicate Spearman correlation coefficients at each time point. Red shaded areas indicate time intervals where a significant correlation (p<0.05; FWE uncorrected) was observed. This analysis was conducted over the entire trial duration with all significant time-points indicated.

**Figure 4.**
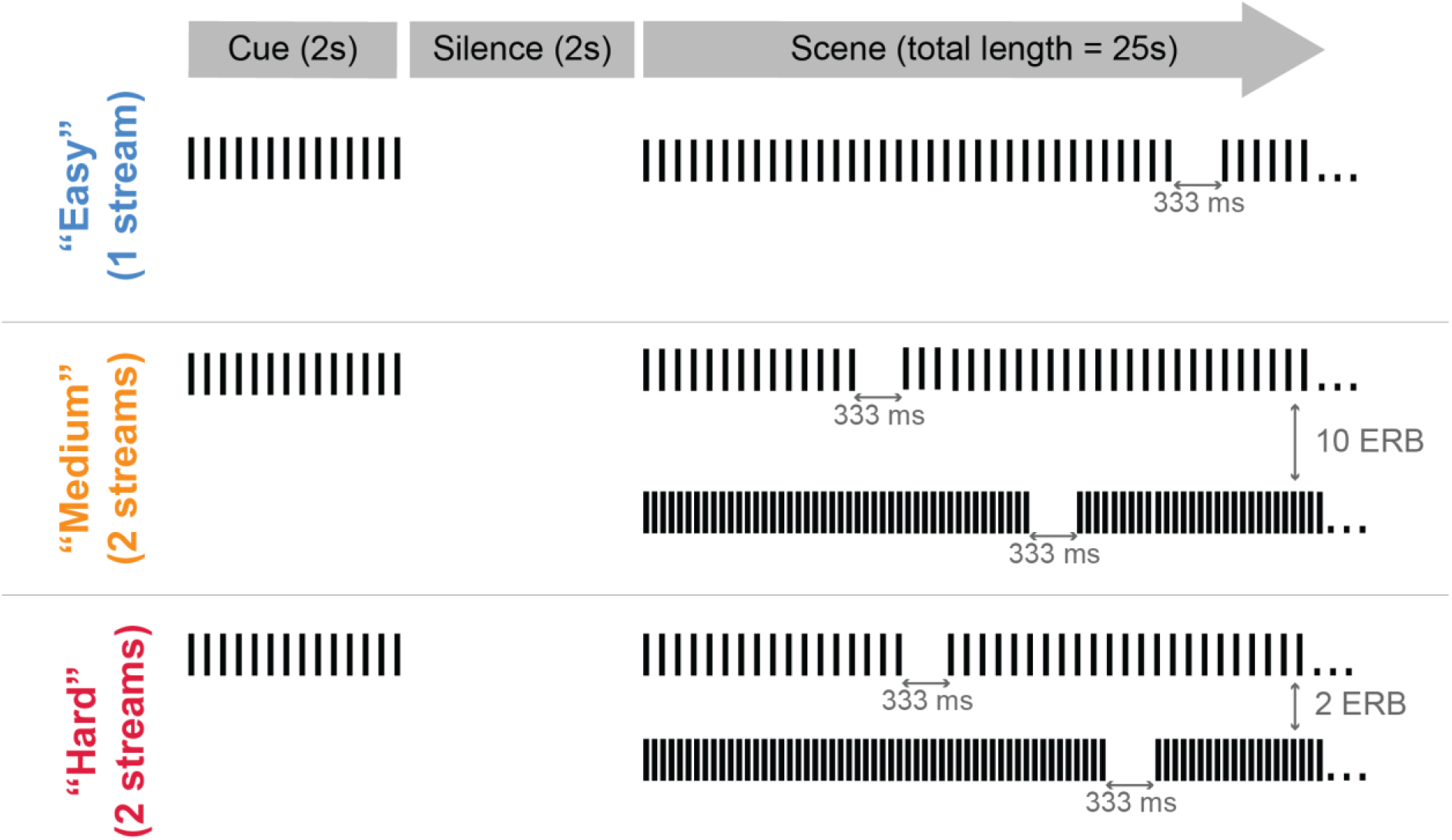
A schematic representation of the stimuli in Experiment 2. Stimuli were similar to those in Experiment 1, with the exception that difficulty was varied by changing the distance between streams. The ‘Easy’ condition consisted of a single stream (identical to that in Experiment 1). The ‘Medium’ condition consistent of 2 concurrent streams separated by 10 ERB. The ‘Hard’ condition consisted of 2 concurrent streams separated by 2 ERB. Other parameters are identical to those in Experiment 1.

**Figure 5.**
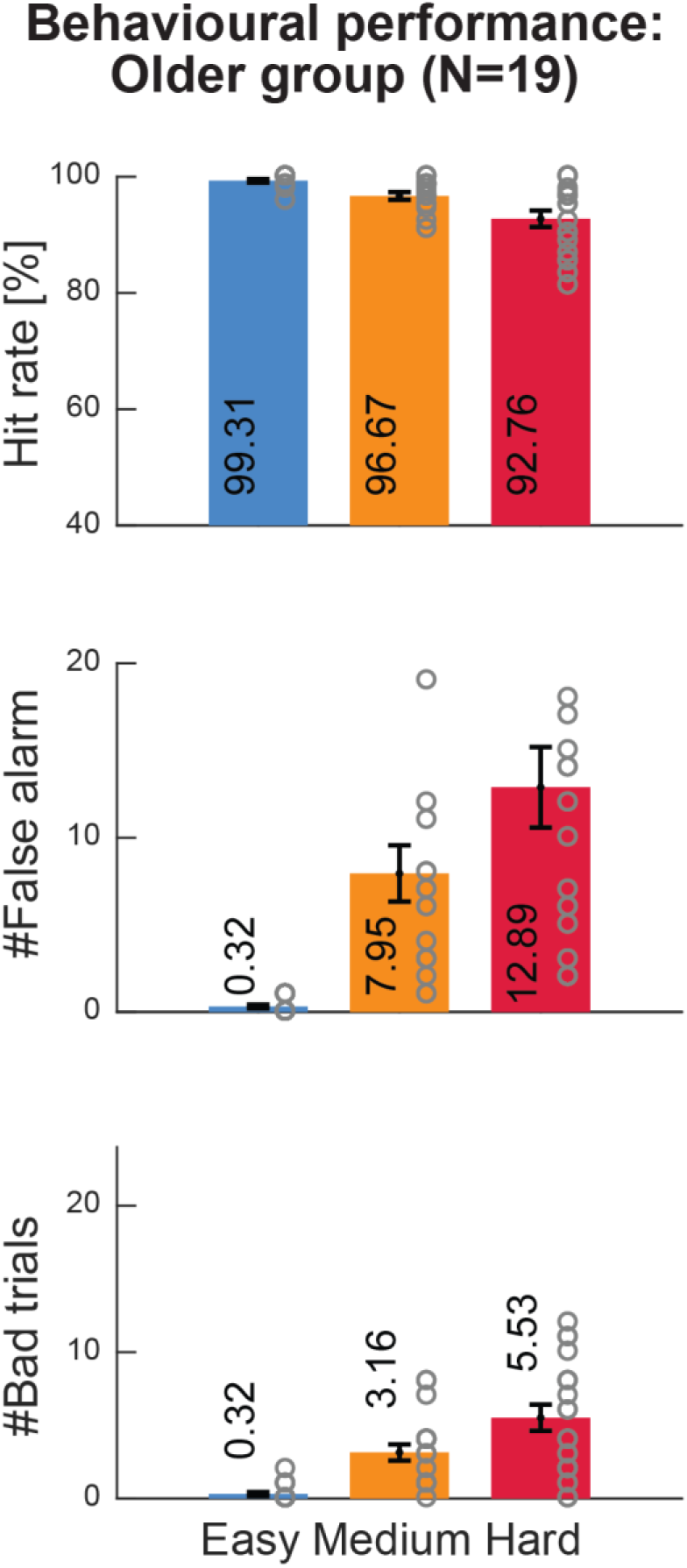
Behavioural performance of the Older group (Experiment 2). Performance measures were: average hit rate, number of false alarms and number of bad trials for retained participants (N=19). Grey circles indicate individual data. All performance measures were significantly modulated by task difficulty. Error bars are ±1 SEM.

To quantify changes to behaviour during the unfolding trial, behavioural data were also analysed over five-time bins of 5s ([0-5]s, [5-10]s, [10-15]s, [15-20]s, [20-25]s). Mean HR and mean #FAs were computed for each condition in each time bin. The data were analysed with a repeated measures ANOVA to investigate main effects of condition and time. The p-value was a priori set to p<0.05. The Greenhouse Geisser correction is used where appropriate.

#### Pupil diameter measurement

An infrared eye-tracking camera (Eyelink 1000 Desktop Mount, SR Research Ltd.) was positioned at a horizontal distance of 65 cm away from the participant. The standard five-point calibration procedure for the Eyelink system was conducted prior to each experimental block and participants were instructed to avoid any head movement after calibration. During the experiment, the eye-tracker continuously tracked gaze position and recorded pupil diameter, focusing binocularly at a sampling rate of 1000 Hz. Participants were instructed to blink naturally during the experiment and encouraged to rest their eyes briefly during inter-trial intervals. Prior to each trial, the eye-tracker automatically checked that the participants’ eyes were open and fixated appropriately; trials would not start unless this was confirmed.

#### Analysis: Pupillometry

As described above, only “successful trials” were included in the pupillometry analysis. To equate the number of trials analysed per condition, the number of trials per condition was set to 12 per participant (this number was determined based on the performance of the worst retained participant on the most difficult condition).

##### Preprocessing

Only the left eye was analysed. To measure the pupil dilation response (PDR) associated with tracking the acoustic stream, the pupil data from each trial were epoched from 0.5 s prior to stream onset to stream offset (25 s post-onset). For each trial, baseline correction was applied by subtracting the mean pupil diameter over the pre-onset interval (0.5-sec pre-onset). The data were smoothed with a 150 ms Hanning window and down-sampled to 20 Hz. Intervals where full or partial eye closure was detected (e.g. during blinks) were automatically treated as missing data and recovered using shape-preserving piecewise cubic interpolation. The blink rate was low overall. In both young and older (see below) participant groups and for all conditions, the average blink rate (defined as the proportion of excluded samples due to eye closure) was approximately 5% (SD = 5%). Blinks were distributed evenly over the trial duration.

For each participant, the pupil diameter was time-domain-averaged across all epochs of each condition to produce a single time series per condition.

##### Time-series statistical analysis

To identify time intervals in which a given pair of conditions exhibited PDR differences, a nonparametric bootstrap-based statistical analysis was used (Efron and Tibshirani, 1994). The difference time series between the conditions was computed for each participant and these time series were subjected to bootstrap re-sampling (1000 iterations). At each time point, differences were deemed significant if the proportion of bootstrap iterations that fell above or below zero was more than 95% (i.e. p<0.05). Any significant differences in the pre-onset interval would be attributable to noise and the largest number of consecutive significant samples pre-onset was used as the threshold for the statistical analysis for the entire epoch.

#### Participant exclusion criteria

Participants with more than 50% of bad trials on the hardest condition (3 streams) were excluded from the main analysis.

### Results

#### Behavioural performance

Figure 2A shows behavioural performance across the full group of participants (N=33). The pattern of performance demonstrates that the task becomes increasingly harder with the addition of distractor streams to the scene (manifested by reduced HR and increased #FA and #of bad trials). This suggests that the paradigm successfully manipulates demands on attentive tracking. Thirteen participants performed poorly on the hardest condition, resulting in an insufficient number of “successful trials”. These participants were excluded from further analysis. The fact that 30% of participants are excluded suggests that the task loads resources to the extent that it may deplete them in a large proportion of participants.

Figure 2B plots the performance of the 20 retained participants (those who had at least 12 successful trials in the hardest condition). A repeated measures ANOVA with condition (1 stream - ‘Easy’, 2 streams - ‘Medium’, 3 streams - ‘Hard’) revealed a main effect of condition on all performance measures (HR, #FA, #bad trials). For HR F(1.464, 27.812) = 36.915, p < .001, for #FA F(1.557, 29.589) = 29.910, p < .001, for #bad trials F(2,38) = 39.181, p < .001. Post-hoc tests (Bonferroni corrected) for HR revealed no significant difference between the ‘Easy’ and ‘Medium’ conditions (p = .065) but did show a significantly reduced HR for ‘Hard’ compared to ‘Easy’, and ‘Medium’ compared to ‘Easy’ trials (p-values < .001). There were significant differences between all conditions for the #FA (p-values ≤.002) and #bad trials (p-values ≤ .026), showing an increase in false alarms and bad trials for conditions with higher numbers of streams.

In addition to quantifying the overall effects, we examined how performance evolved over the duration of the trial by separating the trial into 5s time bins (Figure 3B). A repeated measures ANOVA with condition (‘Easy’, ‘Medium’, ‘Hard’) and time bin (five 5s intervals) revealed a main effect of condition for both measures; for HR F(1.504,28.584) = 33.876, p < .001, for #FA F(1.557, 29.589) = 29.910, p < .001). There was also an interaction between condition and time bin for HR F(8,152) = 3.667, p = .001 and for #FA F(3.996,75.933) = 2.623, p = .041. This was because the difference between the hardest condition and the other two conditions was not fixed but increased partway through the trial. A post hoc repeated measures ANOVA analysis on hit rates in each condition as a function of time-bin revealed no effect for the ‘Easy’ and ‘Medium’ conditions, but a significant effect of time-bin on the “Hard” condition (F(4,76)=8.79 p<.001). An identical result was obtained for analysis of #FA (“Hard” condition F(4,76)=4.08 p=0.05; other two conditions n.s).

#### The Pupil Dilation Response (PDR) as a measure of effort to sustain attention

Figure 3A plots the average pupil diameter data across the 20 listeners as a function of time relative to the pre-onset baseline. Note that the baseline was not taken at a complete resting state but during a brief silent interval (2 seconds) that occurred between the presentation of the cue and the onset of the scene. At this point all conditions are equiprobable.

All three conditions share a similar PDR pattern: Immediately after scene onset (t=0), the pupil diameter rapidly increased and reached a peak within 2 seconds. A significant difference between the PDR to the ‘Easy’ versus ‘Medium’ and ‘Hard’ tracking conditions emerged roughly 1 second after onset. The difference between the ‘Medium’ and ‘Hard’ conditions emerged 2.15 seconds after onset. After the initial peak in the ‘Hard’ condition (at 2 seconds), the pupil diameter continuously climbed to a peak at 4.1 seconds.

Following the initial dilation, the pupil diameter gradually decreased throughout the epoch but in a manner that preserved the differences between the different conditions. The difference between the ‘Medium’ and ‘Easy’ conditions was no longer significant after 14.25 s. However, the PDR to the ‘Hard’ condition remained considerably above the other two conditions throughout the epoch.

Note that the negative pupil diameter values later in the trial reflect the fact that pupil diameter reduced beyond its size during the pre-trial (baseline) period. This likely happens due to the presence of pupil dilation in the pre-trial period, reflecting the anticipation of the onset of the scene (e.g. Bradshaw, 1968; Wierda et al. 2012).

#### Correlation between PDR and behaviour at an individual level

To investigate the relationship between pupil dynamics and behavioural performance on an individual subject level, we correlated within each time bin the HR difference between the ‘Hard’ and ‘Medium’ conditions (Figure 3C) with the corresponding mean PDR difference. The ‘Easy’ condition was excluded from this analysis because it was associated with little behavioural variability across participants, consistent with ceiling performance. Correlation coefficients (Spearman) are plotted in Figure 3D. A significant (Bonferroni corrected), moderate correlation between PDR and HR was observed between 15-20 s after trial onset. This timing corresponded to the time window where the HR and #FA of the ‘Hard’ condition demonstrated increased divergence relative to the ‘Medium’ condition (Figure 3B).

For a more time-sensitive analysis, we also correlated the instantaneous PDR difference between the ‘Hard’ and ‘Medium’ conditions at every time sample (20 Hz) with the mean overall HR difference between these conditions measured for each participant (Figure 3E). Correlation coefficients (Spearman) are plotted as black bars in Figure 3E. Significant time samples (FWE uncorrected) are marked in red. In line with the time-binned analysis, a significant correlation between instantaneous PDR and HR was found between ~12 and ~19 s post-stream onset.

## Experiment 2: Older listeners

Overall, the results from Experiment 1 indicate that pupil dilation is a stable and sensitive measure of effort to sustain attention at the group level and that it is modulated by individual subject performance. This finding makes PDR a potentially useful objective tool for evaluating attentive tracking ability. Specifically, PDR may be instrumental for quantifying deficits in attentive tracking often exhibited by older populations. However, a potential drawback is the known physiological changes to the pupil that occur during healthy ageing; increased demands on accommodation, reduced pupil diameter and slower responses are commonly observed (Bitsios et al., 1996; Guillon et al., 2016; Tekin et al., 2018). Whilst the physiological underpinnings of these effects are not fully clear (Bitsios et al., 1996), they manifest as relative pupil rigidity and may reduce the sensitivity of the PDR as a measure of effort.

In Experiment 2 we used a paradigm similar to that in Experiment 1 to measure attentive tracking capacity in a group of older listeners.

### Methods

#### Participants

Twenty paid participants aged 60 years or older (14 females, average age 70.5, range 63-79) participated in this experiment. Participants were recruited from the U3A (https://www.u3a.org.uk/) and therefore represent a sample of high functioning older individuals. One participant was excluded due to poor behavioural performance (>50% bad trials). All reported no neurological or existing ophthalmological disorders. Several of the participants reported having successfully undergone cataract surgery 2+ years before the present study. Additional inclusion criteria included near normal hearing (see ‘audiometric profile’, below) and normal-range performance on an MCI (mild cognitive impairment) screening test (Addenbrooke’s Cognitive Examination-mobile test). Experimental procedures were approved by the research ethics committee of University College London and written informed consent was obtained from each participant.

##### Audiometric profile

Participants were recruited to this experiment based on evidence of near-normal hearing. This was defined as (air-conducted) pure-tone thresholds of 30 dB HL or better at octave frequencies from 0.25 to 4 kHz in both ears. This range was representative of the frequencies used in our stimuli.

#### Stimuli and Procedure

The stimulus paradigm (Figure 4) was similar to that in Experiment 1 though we made the task somewhat easier. This addressed the concern that the 3-stream task (on which 30% of young participants exhibited low performance) may be too difficult for the older listeners who have previously been demonstrated to be more distractible than young controls (Chadick et al., 2014; Mishra et al., 2014; Petersen et al., 2017). We, therefore, chose to limit the scene to two streams. The difficulty was manipulated by varying the spectral separation between streams. The stimulus conditions here included 1 stream (‘Easy’; identical to experiment one), 2 streams spaced at 10 ERB (‘Medium’) and 2 streams spaced at 2 ERB (‘Hard’). Note that even in the ‘Hard’ condition, the spectral separation is such that streams are sufficiently far apart to limit energetic masking between concurrent sources. Otherwise, stimulus generation, procedure and analysis were identical to those described for Experiment 1. As in Experiment 1, during the practice session participants were allowed to adjust the level at which the stimuli were presented to a comfortable loudness. Older listeners tended to choose a higher level than the participants in Experiment 1.

### Results

#### Behavioural performance

Figure 5 shows the behavioural results in the tracking task. A repeated measures ANOVA with condition (‘Easy’, ‘Medium’, ‘Hard’) as a within-subject factor revealed a main effect for all measures of performance (HR, #FA, #bad trials): for #HR F(1.523, 27.419) = 18.047, p < .001, for #FA F(1.352, 24.344) = 25.418, p < .001, for bad trials F(1.358,24.449) = 25.806, p < .001. Post-hoc tests (Bonferroni corrected) revealed that HR was significantly reduced between each pair of conditions (p ≤ .013). # FA was significantly increased between each pair of conditions (p ≤ .003). #bad trials was significantly increased between each pair of conditions (p ≤ .002).

In addition to quantifying the overall effects, we examined how performance evolved over the duration of the trial by separating the trial into 5s time bins (Figure 6B). A repeated measures ANOVA with condition (‘Easy’, ‘Medium’, ‘Hard’) and time bin (five 5s intervals) revealed a main effect of condition for both HR and #FA measures; for HR F(1.523,27.419) = 18.047, p < .001, for FA F(1.358, 24.449) = 25.806, p < .001. Post-hoc tests revealed that HR and mean #FA were both significantly different between each pair of conditions (p ≤ .009; p ≤ .003 respectively). There was a main effect of time bin for #FA (F(4,72) = 3.128, p = .020) but not HR (F(4,72) = 0.943 p = .444). This effect was driven by an overall smaller number of false alarms in the first bin relative to bin#2 and bin#3.

**Figure 6.**
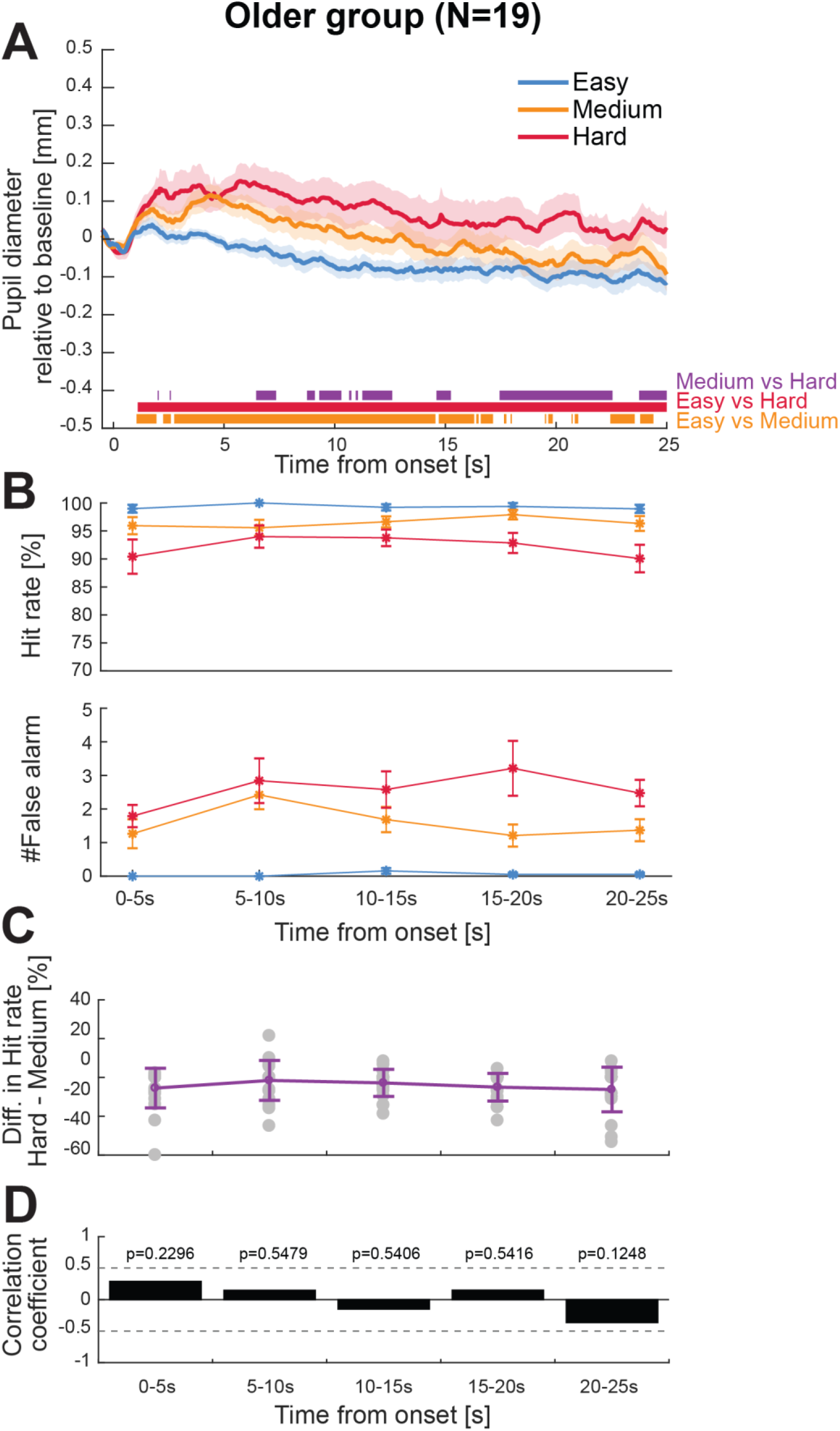
The pupil dilation response reflects effort to sustain-attention. [A] Pupil dilation results from the older group (N=19). The solid lines represent the average pupil diameter as a function of time relative to the baseline (500 ms pre-onset). The shaded area shows ±1 SEM. Colour-coded horizontal lines at graph bottom indicate time intervals where bootstrap statistics confirmed significant differences between each pair of conditions. [B] Time-binned behavioural performance. Error bars are ±1 SEM. [C] Time-binned Hit Rate difference between the ‘Hard’ and ‘Medium’ conditions. Error bars are ±1 standard deviation. Grey dots represent individual data. [D] Correlation between PDR and HR for each time bin. Within each time-bin average PDR difference between the ‘Hard’ and ‘Medium’ conditions is correlated with the corresponding HR difference (as in [C]). Black bars indicate Spearman correlation coefficients at each time bin. No significant correlations were observed.

Hit rates were higher than anticipated (and overall higher than those exhibited by the younger group; though note the task for the younger participants was harder). However, false alarm numbers were equivalent to those exhibited by the younger listeners in Experiment 1, despite the lower difficulty of the task in Experiment 2. This is consistent with an increased propensity for distraction in older listeners.

#### The Pupil Dilation Response (PDR) as a measure of effort to sustain attention

Figure 6A plots the average pupil diameter data across the older listener group (N=19) as a function of time relative to the pre-onset baseline. As for the young group, the data were baselined relative to the silent interval which preceded the scene onset. Consistent with the observations from Experiment 1 (Figure 3A), the older listeners’ pupil response also revealed a stable, positive relationship between the amount of effort required to sustain attention during listening and the pupil diameter.

The PDR to the ‘Hard’ and ‘Medium’ conditions exceeded the PDR to the ‘Easy’ condition from 1.1 s post onset; the PDR to the ‘Hard’ condition also exceeded the PDR to the ‘Medium’ condition from 6.4 s. However, unlike for the young group, we failed to find any systematic relationship between the PDR and individual performance (Figure 6C, D). There could be several reasons for this including factors associated with task difficulty or lack of sufficient pupil reactivity in the older population.

Indeed, it was evident that the PDR exhibited by the older group was overall much smaller and with substantially less across-subject variability than that of the young group. Figure 7A demonstrates that this difference is maintained even when selecting the top performers in each group. Figures 7B, C show that the difference in variability is manifested even when comparing the response to the easiest condition (single stream, identical across groups). This pattern is consistent with the differences between young and older participants observed in the resting state data (Figure 8, below) and is likely to be driven by physiological changes to pupil reactivity.

**Figure 7.**
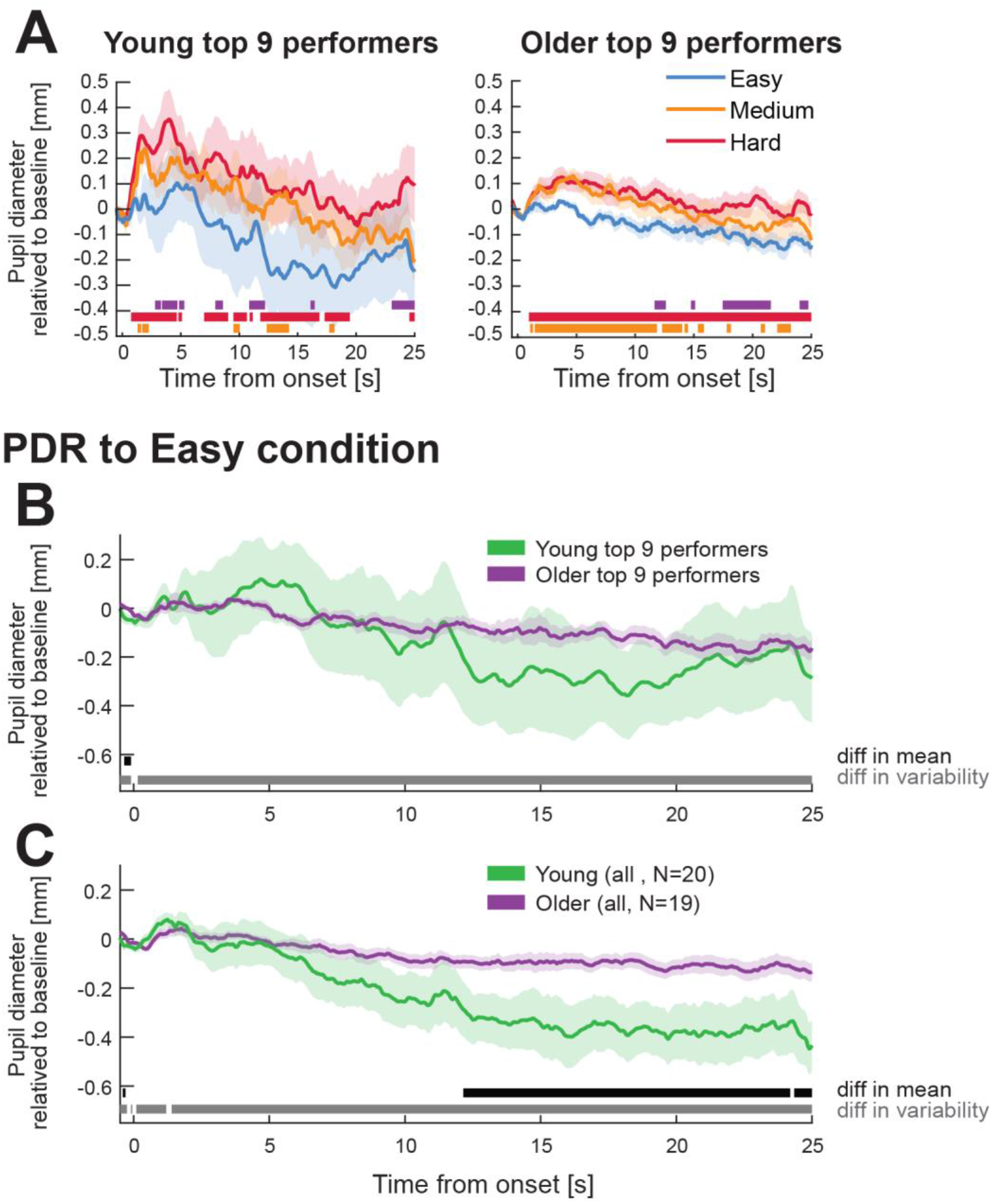
Comparison of the PDR in the young and older groups. [A] Average PDR for the top 9 performers (based on the number of bad trials in the ‘Hard’ condition) for young (left) and older (right) groups. [B] A comparison of the PDR to the ‘Easy’ condition across the top performers in each group. The grey horizontal line shows the significant differences in between-subjects variability (1 standard deviation) and the black line shows the significant differences in the group mean. Despite the fact that this condition was identical across groups, the young group exhibited a substantially larger between-subject variability than the older group. This difference held throughout the epoch. [C] Same as [B] but across all participants in each group. The older listeners exhibited substantially smaller inter-subject variability, consistent with reduced pupil reactivity.

**Figure 8.**
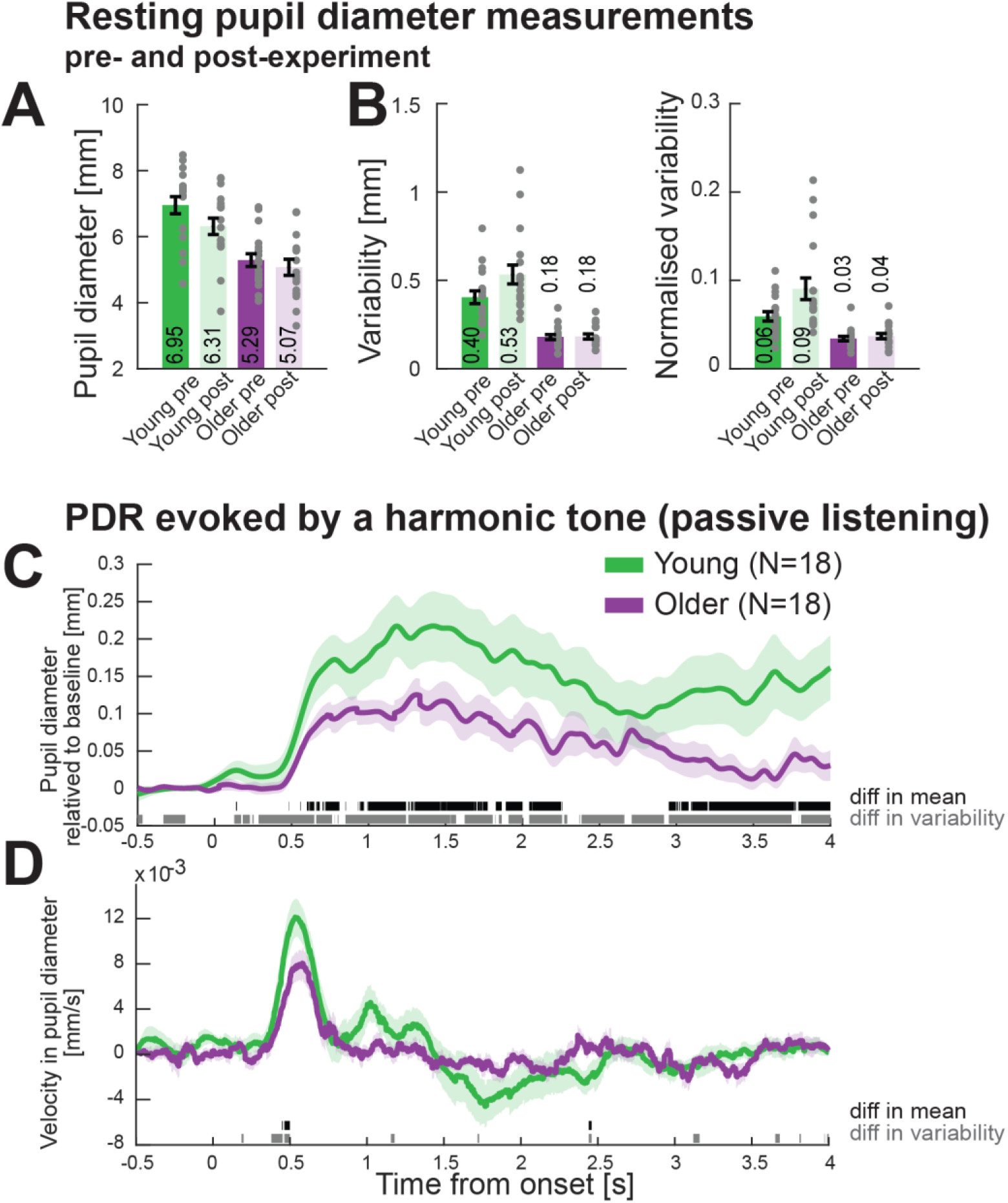
Pupil metrics for the young (N=18) and older (N=18) groups. [A] Median pupil diameter computed over a 30 s ‘resting state’ period, pre- and post-main experiment. [B] Variability (standard deviation) in pupil dimeter over the resting state measurement. The normalised variability is computed by dividing by the mean diameter. The grey circles indicate individual data. Error bars are ±1 SEM. [C] Average pupil diameter as a function of time relative to the onset of a brief harmonic tone. Time intervals where bootstrap statistics show significant differences between the means of the two groups (green and purple lines) are indicated by black horizontal lines. A significant difference in variability (standard deviation) is indicated by grey horizontal lines. [D] The derivative of the pupil data shown in [C] as a measure of the velocity of pupil diameter change. Significant differences are indicated as detailed above.

## Pupil-Metrics: Young vs older listeners

There are known changes to pupil reactivity with age (Bitsios et al., 1996; Guillon et al., 2016; Tekin et al., 2018; Winn et al., 1994). These include a smaller resting state diameter, a reduced dilation range, and slower velocity of dilation. Here we sought to both replicate these measures and include additional measures of reactivity to brief sounds, as previous reports are mostly focused on reactivity to light flashes.

Towards this aim, we conducted a brief series of simple pupil measurements (“pupil metrics”) in the last 18 participants in each group. These measurements were used to derive statistics about pupil size, variability and response time. Participants completed a 30-second resting state measurement (in silence) before and after the main experiment. They also completed an auditory-evoked PDR measurement which included the presentation of thirty 500 ms harmonic tones (f0=200 Hz; 30 harmonics) with an inter-sound interval randomised between 6 and 7 seconds. Participants listened passively to the sounds while pupil responses were recorded. The screen display remained static (identical to that in the main experiment) and participants maintained fixation on a centrally presented black cross.

### Basic Pupil-Metrics

The **resting pupil diameter** was computed as the median value over the 30-second-long resting state trial. **Variability of the pupil diameter** was calculated as one standard deviation over the same period. **Normalised variability** was calculated as variability divided by the corresponding pupil diameter. **Pupil response time** was quantified as the **timing and velocity of the PDR** to the onset of a harmonic tone. PDR onset time was quantified by bootstrap re-sampling over individual subject data in each group and defined as the first time point from which a significant difference from zero (95% of bootstrap iterations above 0) was sustained for at least 150 ms. Velocity was quantified as the peak derivative during the PDR rise time (see Figure 8).

### Results

Figure 8A plots the median pupil diameter over a 30-second “resting state” measurement session before and after the main experiment. A repeated measures ANOVA on pupil size with timing (pre- or post-the main experiment) as a within-subject measure and age group as between-subject measure revealed a main effect of age group (F(1,34)=22.04, p<.0001), confirming the observation that age is associated with a decreased pupil size (Bitsios et al., 1996; Guillon et al., 2016; Piquado et al., 2010; Tekin et al., 2018; Winn et al., 1994). We also observed a main effect of time (F(1, 34)=10.5, p=.003) with no interaction, confirming that, in both groups, pupil diameter was reduced after the main experiment.

Pupil size fluctuated over time even under constant luminance and without any external stimulation or task. An analysis of pupil variability (standard deviation of pupil size over the 30 second interval; Figure 8B) revealed a main effect of age group (F(1,34)=55.3, p<.0001), a main effect of time (F(1,34)=5.5, p=.025) and an interaction between age and time (F(1,34)=5.02, p=.032). Post hoc tests suggested that the source of the interaction was a null effect in the older group (p=.831). Overall, these results reveal a smaller resting state pupil size and smaller variability in pupil size in the older participants (see also Winn et al., 1994). The data also demonstrate a decrease in pupil size after the experimental session in both groups. The younger participants additionally exhibited a decrease in the variability of pupil size after the experimental session. The lack of effect in older people may be due to floor effects.

The decrease in pupil size and variability after the experimental session may be a consequence of the effort to maintain fixation during the experimental session or else associated with cognitive fatigue that is linked to the attentional task. We correlated (Spearman) this change in pupil metrics (both absolute size and variability) with behavioural measures (HR, #FA for all three conditions). In both groups all tests but one (*) were not significant (p>0.109; * change in variability correlated with HR in the ‘Easy’ condition in the young group, r=−0.56, p=0.016). We, therefore, take this result as indicating (at least for the current N) no evidence for a link between a post-session change in pupil dynamics and individual task-related effort.

An additional simple metric of pupil dynamics is the response to a brief sound event during passive listening. Figure 8C plots the harmonic-tone evoked PDR in the young and older groups. Both groups exhibited a PDR after the presentation of the tone; the average pupil diameter over the first 3 seconds following tone onset was significantly above floor as confirmed by a one-sample t-test in each group (young: t(17) = 5.0745, p<.0001; older: t(17) = 5.1331, p<.0001). The onset of the evoked PDR was 0.46 s in the young group and 0.49 s in the older group. A repeated measures bootstrap confirmed that the PDR of the young group was significantly larger than that of the older group from 0.605 s after onset. We compared the velocity of pupil change between the two groups by taking the first derivative of the PDR (Figure 8D). A significant difference between the two groups was observed at ~0.45 s post onset, during the rise time of the pupil dilation response. This pattern replicates parallel observations in the context of the darkness-reflex amplitude (Bitsios et al., 1996; Tekin et al., 2018).

In addition to differences between group means, there was also a significant difference between in-group variability (quantified as the between-subject variance in each group). This difference was present at baseline and maintained throughout the entire epoch.

In sum, older participants exhibited a slower and smaller pupil dilation (pupil diameter change relative to baseline) and smaller inter-subject variability compared to young participants. All in all, both the PDR and the pupil dynamics measures are consistent with reduced reactivity of the pupil musculature in line with previous reports (Piquado et al., 2010; Tekin et al., 2018). However, it is unclear whether the source of these changes is peripheral (Iris physiology) or rather reflects central deficits in the autonomic system. We will return to this point in the discussion.

## Discussion

The present results reveal that in young listeners, pupil dilation provides a robust measure of effort to sustain attention over long durations that are relevant to real-life listening situations. Task difficulty modulated pupil diameter at the group level and revealed modulations of pupil dilation that were correlated with individual subject performance. The demonstration that robust, sustained effects of pupil dilation are measurable even over long trial durations opens the possibility of using pupillometry as an objective measure of sustained attention and for characterizing failure of attention in various populations. We provide evidence that similar effects are also obtainable from older listeners but with important caveats which will be discussed below.

### Behavioural measures of attentive tracking

To provide tight control of both stimulus features and the behavioural task, we used simple artificial acoustic “scenes” that allowed us to isolate the demands associated with attentive tracking from other concurrent perceptual challenges. We showed that performance decreased substantially with the number of elements (concurrent streams) in the scene (Experiment 1) and was also modulated by their spectral proximity (Experiment 2) suggesting that this task is a suitable model with which to capture the challenges of competition for processing resources in crowded acoustic scenes.

Specifically, the task has several key features: 1) To succeed in this task, listeners must continuously monitor the target stream as even momentary distraction may cause them to miss a target, 2) listeners are required to respond to multiple events within the unfolding sequence, providing precise tracking of attention, 3) the task is devoid of memory and semantic confounds commonly associated with speech stimuli, avoiding interactions that may arise as a consequence of the depletion of resources (Mattys and Wiget, 2011; Schmidt et al., 2015). The use of simple sounds (not speech) also circumvents many practical issues including those related to language proficiency, making the paradigm appropriate for a variety of subjects from children to older listeners.

In future work, the stimuli can be made increasingly complex by varying scene size, source trajectories (e.g. introducing frequency modulation), spatial extent, etc. Due to their narrowband nature, the signals can also be easily adjusted to fit the hearing profile of the individual tested.

### Pupil measures in young listeners track effort to sustain attention

We found that challenging stream tracking conditions were accompanied by large, sustained pupil dilation that mirrored behavioural performance on the group and individual level. Importantly, this sustained pupil dilation cannot be explained by day-dreaming (Unsworth and Robison, 2016) or task disengagement (Hopstaken et al., 2015), because only successful trials were included in the pupillometry analysis.

Despite using time-constant stimulus parameters the behavioural data indicated that task difficulty was not stable but increased partway through the trial. Statistical analysis showed that this particularly affected the hardest condition where the hit rate decreased substantially above the easier conditions from about 10 seconds onwards. The same pattern was present in the pupillometry data. Notably, it was around this time that significant individual-level correlations between pupil diameter and performance were observed. These effects are consistent with multiple observations that the ability to sustain attention deteriorates with time-on-task (Fortenbaugh et al., 2017; Thomson et al., 2015) and is hypothesized to reflect weakened control of cognitive resources (Berry et al., 2017; Esterman et al., 2014; Pattyn et al., 2008; Sarter and Paolone, 2011; Thomson et al., 2015). Indeed, that the false positive rate peaked mid-trial, reflecting the fact that participants were increasingly unable to resist distraction from the non-target streams, is in line with previous proposals that reduced resource control is associated with impaired distractor filtering (Sarter et al., 2001).

Due to our policy of analysing only successful trials, we had to exclude 30% of the young participants who failed to achieve a sufficient number of trials for analysis. That about a third of our cohort failed on the hardest condition suggests that resources were indeed likely exhausted by the task. Whether there are any cognitive markers which might differentiate those who succeeded from those who failed is an interesting question for future work.

### Is pupillometry in older listeners a useful objective measure?

Ageing is associated with loss of function within the peripheral auditory system that leads to a broad range of auditory processing impairments. In addition, normal ageing is associated with various deficits of cognitive, executive and sustained-attention function that have expansive perceptual consequences across sensory modalities. In the context of the hearing, these deficits may have wide-ranging implications for listening in crowded environments, such as the ability to attend to a relevant sound source and avoid distraction by concurrent sounds. Routine audiological assessments are not sensitive to these impairments, resulting in sub-optimal understanding and management of these conditions. Pupillometry may be a promising tool to quantify such impairments as it is cheap, portable and non-invasive.

We observed clear and robust effects of task difficulty on pupil diameter in our group of normal hearing, high performing older individuals. These effects were sustained over the trial duration and paralleled group-level behavioural performance (Figure 6). However, they also differed from those observed by the young group in several important respects:

Firstly, unlike in the young group, we did not see any correlation with individual performance. This may be because the task was too easy. Although we decided on the present task setting based on pilot experiments, the resulting performance was better than expected. Future work should adjust the difficulty to each listener independently.

Secondly, the pupil data from older participants exhibited substantially smaller variability across participants and time. This cannot be due to the overall higher performance levels observed on this task because similar effects were also present in the “pupil metrics” data (Figure 8) that were acquired under passive listening conditions. These effects are in line with known age-related changes to ocular physiology. One of the major changes even with no clinical sign of disease is increased rigidity of the pupil (senile miosis; Meller, 1904) resulting in overall decreases in pupil size, range of pupil dilation, variability and response speed (Bitsios et al., 1996; Tekin et al., 2018).

Previous work raised the concern that the restricted range of the pupil in older listeners may limit the ability to see small, cognitive-state mediated changes to pupil size (Piquado et al., 2010; Van Gerven et al., 2004). However, here we observed significant sustained effects despite quite small behavioural differences between conditions, suggesting that pupillometry can be a sensitive measure of effort in this population.

There are several potential explanations for the effects we observe in the older cohort. One possibility is that the temporal variability in pupil size present in young listeners may reflect physiological noise. The lack of such variability in the older group may thus be taken as an advantage. However, it is increasingly understood that instantaneous fluctuations in pupil size reflect momentary changes in perceptual state that contribute in important ways to behavioural variability (Allen et al., 2016; Fontanini and Katz, 2008; Kelly et al., 2008). The reduced range of the pupil in older populations may make us blind to many of these effects.

Another, not mutually exclusive, possibility relates to the mechanisms that support pupil dynamics. As we discuss further below, both sympathetic and parasympathetic systems can affect pupil dilation. It is feasible that the general rigidity of the pupil in older participants may be related to a reduction in sympathetic activity (Bitsios et al., 1996), whilst the effects observed during attentive tracking are produced by the relatively preserved parasympathetic activity.

### Neuromodulator effects on sustained attention

Mounting evidence from electrophysiology in animal models has revealed a strong correlation between pupil-size dynamics and activity of NE (Joshi et al., 2016; Phillips et al., 2000; Rajkowski et al., 1993) and ACh expressing neurons (Reimer et al., 2016; Zaborszky et al., 2015). Pupil size is modulated by the balance between dilator and sphincter muscles in the iris. The dilator muscle is innervated by the sympathetic system which acts by releasing NE, and the sphincter muscle is innervated by the parasympathetic system for which ACh is the major neurotransmitter (Loewenfeld and Lowenstein, 1993; Steinhauer and Hakerem, 1992; Steinhauer et al., 2004). ACh exerts an inhibitory effect in the oculomotor nucleus of the brain stem leading to relaxation of the sphincter muscles, and therefore also to pupil dilation. Consequently, increased release of NE and ACh both contribute to pupil dilation (Larsen and Waters, 2018), however, whether the effects are independent or also synergistic remains unknown.

NE and ACh are hypothesized to play key roles in supporting cognitive effort and executive control (Aston-Jones and Cohen, 2005; Botvinick et al., 2001; Sarter et al., 2006; Steinhauer et al., 2004). Specifically, a large body of work has linked NE release to increased arousal (see review Berridge and Waterhouse, 2003) and sustained attention (e.g., Aston-Jones et al., 1994; Carli et al., 1983; Sara 2009). ACh has been associated amongst other things with activation in the anterior attention system (which underlies effortful, top-down control of goal-directed behaviour; Petersen and Posner, 2012) and is hypothesized to play a role in controlling distraction (Berry et al., 2014; Demeter and Sarter, 2013; Kim et al., 2017; Sarter et al., 2006; Himmelheber et al, 2000). In the context of the present task, it is possible that the observed pupil dilation effects reflect NE-mediated heightened vigilance as well as ACh-mediated processes linked to the need to maintain focus on the target sequence and avoid distraction from the concurrent, non-target auditory streams.

Based on the pupillary response pattern observed in their experiments, Bitsios et al.(1996; see also Tekin et al., 2018) argued that the altered pupil dynamics commonly observed in ageing subjects and also replicated for the present cohort (Figure 8) are of a central origin and predominantly driven by weakened signalling from the sympathetic system.

It is therefore tempting to postulate that the reduced variability in pupil diameter observed here in older listeners (Experiment 2) may reflect the decline in NE-mediated pupil dilation whilst the preserved effect of attention on average pupil size may be specifically driven by ACh-linked pupil dynamics. This is also consistent with a key role for ACh in supporting attentive listening by suppressing distractors – a main feature of the present task (Berry et al., 2014; Demeter and Sarter, 2013; Himmelheber et al., 2000; Kim et al., 2017; Sarter et al., 2006). Future work with more sensitive techniques in animal models or pharmacological manipulations in humans (see also Steinhauer et al., 2004; Wang et al., 2016) is needed to tease apart the contribution of ACh and NE to attentive listening.

## Acknowledgements

Supported by funding from the BRC and the EC (Horizon2020 grant to MC). Initial pilot experiments were performed by Youna Cho. We are grateful to Mathilde Le Gal de Kerangal for helping recruit and screen the older listeners in Experiment 2.

